# Para-allopatry in hybridizing fire-bellied toads (*Bombina bombina and B. variegata*): inference from transcriptome-wide coalescence analyses

**DOI:** 10.1101/030056

**Authors:** Beate Nürnberger, Konrad Lohse, Anna Fijarczyk, Jacek M. Szymura, Mark L. Blaxter

## Abstract

Ancient origins, profound ecological divergence and extensive hybridization make the fire-bellied toads *Bombina bombina* and *B. variegata* (Anura: Bombinatoridae) an intriguing test case of ecological speciation. Narrow *Bombina* hybrid zones erect barriers to neutral introgression whose strength has been estimated previously. We test this prediction by inferring the rate of gene exchange between pure populations on either side of the intensively studied Kraków transect. We developed a software pipeline to extract high confidence sets of orthologous genes from *de novo* transcriptome assemblies, fitted a range of divergence models to these data and assessed their relative support with analytic likelihoods calculations. There was clear evidence for post-divergence gene flow, but, as expected, no perceptible signal of recent introgression via the nearby hybrid zone. The analysis of two additional *Bombina* taxa (*B. v. scabra* and *B. orientalis*) validated our parameter estimates against a larger set of prior expectations. Despite substantial cumulative introgression over millions of years, adaptive divergence of the hybridizing taxa is essentially unaffected by their lack of reproductive isolation. Extended distribution ranges also buffer them against small-scale environmental perturbations that have been shown to reverse the speciation process in other, more recent ecotypes.

## Introduction

The central role of environmental heterogeneity in driving the evolution of novel ecotypes has been impressively illustrated in the rapidly accumulating literature on ecological speciation. In particular, numerous studies have demonstrated how divergent adaptation can lead to partial reproductive isolation by causing assortative mating within ecotypes, reduced performance of hybrids or both (*e.g.* Hatfield and Schluter 1999; Rundle et al. 2000; Via et al. 2000; Jiggins et al. 2001; Linn et al. 2003; Bearhop et al. 2005; Ramsey et al. 2003; Huber et al. 2007; see Nosil 2012 for a detailed treatment). Such barriers to gene flow can arise even on a historical time scale (Hendry et al. 2007). If this momentum can be maintained then, intriguingly, ecological speciation may run its course relatively swiftly.

The hypothesis of rapid ecological speciation appears to be contradicted by the millions of years of divergence that are typically required before substantial pre- and/or postzygotic isolation is observed in experimental settings (Coyne and Orr 1997; Sasa et al. 1998; Price and Bouvier 2002; Bolnick and Near 2005). interbreeding in nature may cease much earlier, if divergent adaptation includes novel habitat preferences and/or shifts in breeding/flowering time (Schemske 2010). But in the absence of irreversible barriers to reproduction, the new ecotypes are reliant on the particular environmental conditions under which they evolved (Boughman 2013). Subsequent change may increase gene flow levels so that phenotypic divergence is eroded again (Taylor et al. 2006; Seehausen et al. 2008; Vonlanthen et al. 2012; Grant and Grant 2014) or it may create periods of allopatry that facilitate the accumulation of habitat-independent gene flow barriers (Butlin et al. 2008; Bierne et al. 2013). In that case, speciation is initiated by ecological divergence but may be completed by a variety of processes. Much work has been done on recently evolved ecotypes (Faria et al. 2014), many of which will not persist long enough to become reproductively isolated (Futuyma 1987; Dynesius and Jansson 2014). It is therefore of interest to determine the level of gene exchange between taxa with a long history of ecological divergence in allo-and parapatry.

Here, we report on one such taxon pair at a relatively advanced stage of the speciation process and estimate the rate of gene flow between two pure populations on a either side of a current hybrid zone. This long term average reflects introgression past genetic barriers that arise from ecological divergence and any habitat-independent incompatibilities. We discuss our results in the context of theory about genetic architectures that produce more or less efficient barriers to gene flow.

The European fire-bellied toads, *Bombina bombina* and *B. variegata,* are clearly defined taxa that differ profoundly in a large number of traits, including features of morphology, anatomy, life history and the mating system (see Szymura 1993 for an overview). Many of these differences reflect the distinct ecological niches that the toads occupy. *B. bombina* reproduces in semi-permanent lowland ponds, whereas *B. variegata* uses ephemeral breeding sites in more mountainous terrain. These aquatic habitats place opposing demands on the tadpoles: cryptic behavior to avoid invertebrate predators *versus* rapid growth and development as desiccation looms (Rafińska 1991; Werner and Anholt 1993). Morphological and behavioral differences between the tadpoles (Vorndran et al. 2002) match established patterns in specialized anuran species along the aquatic permanence gradient (*e.g.* Relyea 2001). Many differences between the adults are also likely adaptations, for example of *B. variegata* to an ever-shifting distribution of breeding sites in the landscape and of *B. bombina* to the large mating aggregations that form in ponds. Such simultaneous selection on multiple traits (multifarious selection) may generate particularly efficient barriers to gene flow (Rice and Hostert 1993; Nosil et al. 2009).

*B. bombina* and *B. variegata* diverged several million years ago (Pabijan et al. 2013) and have experienced repeated cycles of allo- and parapatry (Fijarczyk et al. 2011). Present day hybrid zones formed after the last glacial maximum (less than 10,000 years ago; Szymura 1993, 1996). They are typically 2 - 9 km wide and located along the edge of mountainous regions (Fig. 1). Recombinant marker genotypes (< 10 diagnostic loci) predominate in intensively sampled transects in Poland, Hungary, Romania and Ukraine (Szymura and Barton 1991; Gollmann 1987; Vines et al. 2003; Yanchukov et al. 2006) and make up about a quarter of individuals in the Peŝćenica (Croatia) transect (MacCallum et al. 1998). We expect that all individuals in a hybrid zone are recombinant to some degree. The temporal stability of phenotypic clines at 50 and 70 year sampling intervals (Szymura and Barton 1991; Yanchukov et al. 2006) proves that natural selection counteracts the erosion of taxon differences. Direct evidence for habitat-independent natural selection comes from increased mortality of hybrid embryos under uniform conditions (Szymura and Barton 1986; Kruuk et al. 1999b).

**Figure 1.**
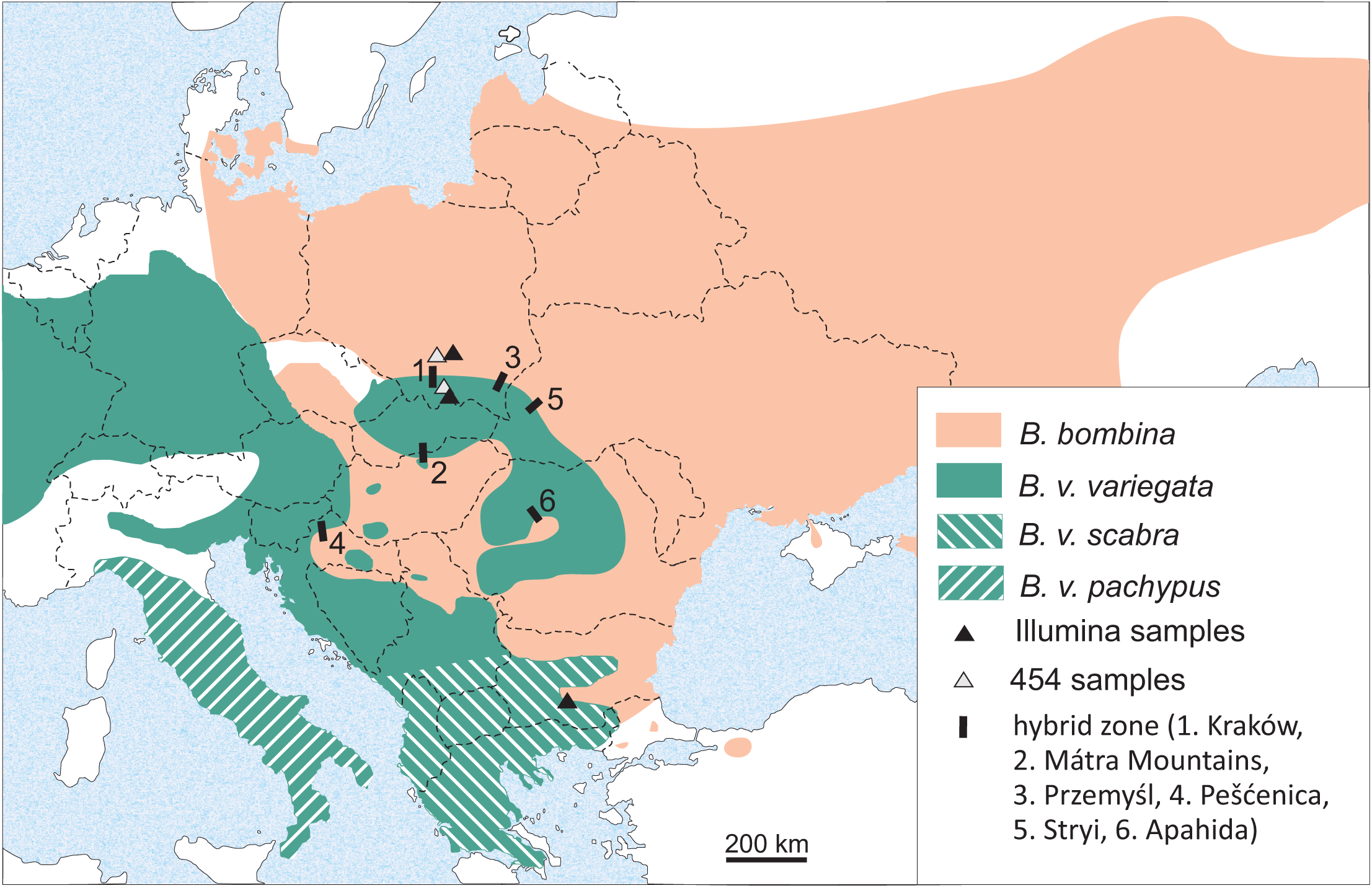
Distribution of *B. bombina* and *B. variegata* in Central and Eastern Europe. Included are the sampling locations (triangles) and the approximate locations of intensively studied hybrid zones. Dashed lines represent national boundaries.

Classic cline theory links the patterns seen in nature to the three main determinants of hybrid zones: gene flow, selection and recombination. It predicts that the observed significant linkage disequilibria among independently segregating markers (Nürnberger et al. 2003) result from the influx of pure genotypes into the hybrid zone center and increase the ‘effective’ selection pressure on each one of the selected loci (Szymura and Barton 1986; Barton and Gale 1993; Kruuk et al. 1999a). The associated allele frequency clines thus become steeper (Slatkin 1973) which further strengthens linkage disequilibria and effective selection. This positive feedback (Barton 1983) can explain the sharp step in allele frequency in the center of all clinal *Bombina* transects (Szymura and Barton 1991; Yanchukov et al. 2006). It helps to maintain adaptive taxon differences and also generates a barrier to gene flow for neutral variants (Nagylaki 1976; Barton 1986).

The strength of this barrier reflects the overall level of reproductive isolation and can be estimated from the shape of clines at marker loci (Szymura and Barton 1986; Barton and Gale 1993). The two Polish transects near Kraków and Przemyśl (width: ~ 6 km) provide a robust estimate of this barrier into the *B. variegata* gene pool (Szymura and Barton 1991). Under a diffusion model of dispersal and based on an estimated dispersal range of ~ 1 km per generation (Szymura and Barton 1991), this barrier is equivalent to 51 km of unimpeded habitat. Assuming a generation time of three years (Szymura 1988), it would delay the introgression of neutral *B. bombina* alleles by about 8000 years (Barton and Gale 1993). The estimated barrier strength also implies that the mean fitness of hybrid populations in the zone center is reduced by 42% relative to that of pure populations (Szymura and Barton 1991). Mitochondrial genomes might be expected to introgress more freely (Toews and Brelsford 2012), but this is not observed in *Bombina:* concordant mtDNA and nuclear marker clines together with very weak cytonuclear disequilibria suggest that mtDNA may itself be subject to selection in hybrids (Szymura et al. 1985; Yanchukov et al. 2006; Hofman and Szymura 2007; Fijarczyk et al. 2011; but see Vörös et al. 2006).

Thanks to the genome-wide linkage disequilibria in the hybrid zone, it was possible to draw all of these inferences from the small number of available diagnostic markers. Direct estimates of gene exchange between pure populations, however, require genome-wide genetic resources. Their development has so far been hampered by the ‘big’ genomes (Dufresne and Jeffery 2011) of *B. bombina* and *B. variegata;* for both taxa, C-value estimates just over the threshold of 10 pg (~ 10 Gbp) have been obtained (Gregory 2015). There is, however, no indication of polyploidy (Olmo et al. 1982). The two sequenced anuran genomes (*Silurana (Xenopus) tropicalis,* Hellsten et al. 2010; *Nanorana parkeri,* Sun et al. 2015) are too distant from *Bombina* (MRCA about 250 Mya) to be useful. We therefore chose *de novo* transcriptome assembly as a cost-effective method to generate reduced representation genomic data (Davey et al. 2011).

Evolutionary or population genomic analyses of next generation sequence data take a locus-centered view (Cahais et al. 2012) and require the definition of orthologous sequences across samples. With RNA-seq data, the task of ortholog definition is non-trivial, because variants of a single locus (splice isoforms, alleles) are recovered along with sequences from recent paralogs. The identification of orthologs across samples cannot succeed unless paralogous sequences from each sample are included in the search, but the result is ambiguous if splice isoforms are clustered as well. Similarly, the discovery and genotyping of single nucleotide polymorphisms (SNP) based on read mapping runs the risk of either conflating variation from several loci (false positives) or failing to detect allelic variants across different isoforms (false negatives, De Wit et al. 2015).

These issues have been variously addressed in recent *de novo* transcriptome assemblies. With the help of a reference genome, paralogs can be identified by phylogenetic methods (*e.g.* Heger and Ponting 2007; Osborne et al. 2013). Alternatively, variant call sets have been screened for suspect genotype distributions in order to remove SNPs that are likely derived from more than one locus (*e.g.* Gayral et al. 2013; Harris et al. 2015). In the absence of a reference genome and starting from either one or two samples per taxon, we devised criteria to identify putative paralogs within transcriptome assemblies. These were included in a clustering step across taxa to generate a high-confidence ortholog gene set for which SNPs were called. Each orthogroup contained sequences from four *Bombina* taxa with a fully resolved phylogeny based on complete mitochondrial genomes (Zheng et al. 2009; Pabijan et al. 2013): the hybridizing pair *B. bombina* and *B. v. variegata* as well as *B. v. scabra,* a subspecies from the Southern Balkans (Fig. 1), and *B. orientalis* (East Asia), the sister taxon of the European lineages.

To quantify nuclear divergence and introgression, we used a recently developed coalescent approach (Lohse et al. 2015) that fits models of divergence with or without gene flow and assesses their support via analytically computed likelihoods. This inference is extremely efficient compared to multilocus methods that require computationally intensive Markov chain Monte Carlo sampling (Hey and Nielsen 2004) or simulations (*e.g.* Shafer et al. 2015). Models were fit by two approaches: either using the site frequency spectrum (SFS) or multilocus data summarized by the blockwise SFS, *i.e.* the full configurations of mutations in blocks of sequence. With these coalescent analyses, we address the following questions: (i) Does the timing and order of divergence between the four *Bombina* lineages match that previously inferred from mitochondrial data?, (ii) Is there evidence for post-divergence gene flow between the hybridizing taxa away from the hybrid zone and, if so, how does it compare to the estimate between the *B. variegata* subspecies (genetic barrier to gene flow *vs*. greater spatial separation)? and (iii) Is there any evidence for a recent burst of introgression as a result of the current hybrid zone?.

## Methods

### Sample origins

Roche 454 sequence data were generated from two females: *B. bombina (Bb),* Kozłów, Poland (50°29’N, 20°02’E); *B. v. variegata (Bvv),* Mogielica, Poland (49°39’N, 20°16’E). Illumina data came from four individuals: *B. bombina,* male, Sędziejowice, Poland (50°34’N, 20°39’E); *B. v. variegata,* male, Biała Woda, near Szczawnica, Poland (49°24’N, 20°34’E); *B. v. scabra (Bvs),* female, Rhodopes, Bulgaria; *B. orientalis (Bo),* female, Northern clade, Korea. The European sampling locations are indicated in Figure 1. The *B. bombina* and *B. v. variegata* sample sites are on either side of the Kraków transect (Szymura and Barton 1986, 1991) and their distances to the cline center are approximately 61 km (Bb, Kozłów), 72 km (*Bb*, Sędziejowice), 36 km (*Bvv*, Mogielica) and 61 km (*Bvv*, Biała Woda).

### RNA extraction, library preparation and sequencing

For Roche 454 sequencing, total RNA was extracted from liver, using RNAzol®RT, and mRNA was isolated using Dynabeads Oligo(dT)_25_. The RevertAidTM Premium First Strand cDNA Synthesis Kit was used to synthesize the first cDNA strand with a primer complementary to the poly(A) tail with disrupted sequence: 5’-TTTTTCTTTTTTCTTTTTTV-3’. The RNA in the RNA:DNA hybrid was cleaved with RNAse H and the second strand synthesized using DNA polymerase I and T4 DNA polymerase. No additional PCR reactions were performed and the cDNA library was not normalized. Two cDNA libraries were sequenced independently (in half a FLX plate each) on a 454 GS FLX Titanium system (454 Life Sciences, Roche) at the Functional Genomics Center in Zürich (Switzerland).

For Illumina RNA-seq, frozen samples were thawed and transferred into 1 ml Trizol, homogenized in a TissueLyser and passed five times through a 0.9 mm (20 gauge) needle attached to a syringe (Gayral et al. 2013), followed by 5 min incubation at RT and centrifugation for 1 min at 14,000 rpm. The samples were Chloroform extracted once and then mixed with 200 μl of 70% Ethanol. Each sample was divided into 2 - 3 aliquots, each of which was added to a Qiagen miRNA spin column and processed according the manufacturer’s instructions. cDNA library preparation and paired-end sequencing on an Illumina HiSeq instrument were carried out by Edinburgh Genomics.

### Transcriptome assemblies

*Roche 454 data -* Reads were cleaned by removing residues of poly(A) tails with SnoWhite v.1.1.4 (Barker et al. 2010). Reads with low-complexity regions (including stretches of di- and trinucleotide repeats which could inflict spurious assemblies) and exact read duplicates were also removed. Low quality ends were trimmed with PRINSEQ-lite 0.17 (Schmieder and Edwards 2011). Reads which showed significant sequence identity (using BLASTN) to rRNA sequences of *Silurana tropicalis* were discarded.

*De novo* transcriptome assemblies of *B. bombina* and *B. variegata* were conducted separately for each species. We took a combinatorial approach to optimize each assembly (Kumar and Blaxter 2010). Reads were first assembled using Newbler (v2.6, Roche) and Mira v. 3 (Chevreux et al. 2004) independently, after which both assemblies were co-assembled with CAP3 (Huang and Madan 1999). For Mira and Newbler we used settings recommended for *de novo* transcriptome assembly from reads produced by the 454 platform (Mira: no clipping of poly(A) stretches). For CAP3, we used an overlap length cutoff equal to 80, an overlap percent identity cutoff equal to 95, and switched off the clipping option. To validate the assembly, reads were mapped back to assembled contigs using BLAT (Kent 2002), retaining only contigs with reads that mapped over at least 95% of their length (discarded contigs: 11% of *B. bombina* and 14% of *B. variegata*).

*Illumina data* - Adapter removal was carried out with Cutadapt (Martin 2011) as implemented in the wrapper Trim Galore! v. 0.3.7 (http://www.bioinformatics.babraham.ac.uk/projects/trim_galore/) and quality trimming was performed with Trimmomatic (Bolger et al. 2014), clipping any sequence downstream of four consecutive bases with mean quality below 25. Only sequences with a minimum length of 50 nucleotides (nt) were retained (Table 1).

**Table 1.** Data summary. Listed are the read counts before and after quality control (QC) and assembly statistics for the Illumina and Roche 454 datasets (assembly software in parentheses). Abbr. = taxon abbreviation. Total assembled bases (Trinity) are computed from the longest isoform per component. Only the contigs with read support are listed.

| Taxon | Abbr | Illumina (Trinity) |  |  |  |  |  | Roche 454 (Newbler/Mira/CAP3) |  |  |  |
| --- | --- | --- | --- | --- | --- | --- | --- | --- | --- | --- | --- |
|  |  | raw<br>sequence<br>pairs (x 10<sup>6</sup>) | reads<br>pairs<br>after QC<br>(x 10<sup>6</sup>) | single<br>reads<br>after QC<br>(x 10<sup>6</sup>) | components<br>("genes") | contigs | total<br>assembled<br>bases<br>(x 10<sup>6</sup>) | raw reads | reads<br>after QC | contigs | total<br>assembled<br>bases<br>(x 10<sup>6</sup>) |
| *B. v.*<br>*variegata* | *Bvv* | 55.3 | 42.3 | 7.7 | 67,300 | 80,878 | 47.4 | 636,551 | 611,417 | 22,009 | 15.9 |
| *B. v.*<br>*scabra* | *Bvs* | 34.4 | 25.9 | 5.2 | 53,909 | 62,920 | 36.9 |  |  |  |  |
| *B.*<br>*bombina* | *Bb* | 47.1 | 35.9 | 6.9 | 61,724 | 73,131 | 44.5 | 691,857 | 667,125 | 28,827 | 19.8 |
| *B.*<br>*orientalis* | *Bo* | 46.9 | 35.3 | 6.9 | 52,484 | 62,261 | 35.7 |  |  |  |  |

*De novo* assembly was carried out with Trinity (v. 2014-04-13p1, Grabherr et al. 2011) in paired-end mode with the addition of all retained single reads. Default settings were chosen except for the minimum K-mer count (2), the maximum expected length between fragment pairs (400) and the minimum percent identity required for the merging of paths during transcript reduction (98). Assemblies were searched for contaminants by BLAST analysis (megablast) against the NCBI nucleotide database and contigs with close to full length matches (ID > 0.90) to Bacteria and Archaea were removed (37 - 219 contigs per taxon). For each assembly, a BLASTX comparison to the *Silurana (Xenopus) tropicalis* proteome (ftp.xenbase.org/pub/Genomics/JGI/Xentr7.1/Xentr7_2_Stable_Protein.fa.gz, last accessed 16 Sept 2014) was carried out with expect-value cutoff = 0.01. Read support for each contig was quantified with RSEM (Li and Dewey 2011) using a Trinity script. All contigs with fpkm (fragments per kilobase per million reads) = 0 were removed from the assemblies (2514 - 4004 contigs per taxon). Open reading frame (ORF) predictions were computed with TransDecoder which is bundled with Trinity. Finally, assembly completeness was assessed with CEGMA (Parra et al. 2007).

### Reference transcriptomes

For each taxon, we aimed to define a reference transcriptome that comprised all assembled gene copies (including paralogs) but excluded splice isoforms. Details of the analyses are presented in the Supporting Information and summarized here. Sequence identities (ID) were computed for pairwise nucleotide alignments within Trinity components. In the first instance, all contig pairs with overall sequence identity below 0.98 were designated as paralogs. In the case of markedly heterogeneous ID scores along the alignment, we distinguished poorly aligned tracts due to structural variation near alignment termini from those due to the alternate use of duplicated exons. The latter case represents paralogy below the level of the gene (*cf.* Gabaldón and Koonin 2013) and contigs were designated accordingly, even if the mean ID was greater or equal 0.98. In the case of *B. bombina* and *B. v. variegata,* additional pairwise comparisons were carried out, using BLASTN, of the Roche 454 contigs to the Trinity contigs. Paralogs were identified as before. For each taxon, the reference consisted of one contig per Trinity component (with maximum ORF length), all associated paralogs and, if available, all Roche 454 contigs (separated into paralogs and non-paralogs).

### Orthologue identification and alignment

We used OrthoMCL (Li et al. 2003) with default parameters to cluster predicted peptide sequences from all four reference transcriptomes (Fig. 2). The complete proteome of S. *tropicalis* was added as a ‘backbone’ for clustering and to provide additional information on multi-gene families. From the initial clustering (inflation value = 1.5), we retained only those groups that comprised the following: (1) at least one contig from each *Bombina* taxon, (2) no paralog pairs from any *Bombina* taxon and (3) no more than three S. *tropicalis* contigs (see Fig. 2b for examples). From this subset of clusters we retained only those that were robust to changes in the inflation value (range: 0.5 - 5.0) in order to counter common errors of graph-based methods such as OrthoMCL, *i.e.* over-clustering and incorrect handling of domain recombination (Kristensen et al. 2011). Finally, pairwise peptide alignments between *Bombina* taxa were computed within robust orthogroups. Note that we apply stringent criteria to obtain a set of robust orthogroups rather than an accurate list of paralogs. False positives in that list are therefore of no concern.

**Figure 2.**
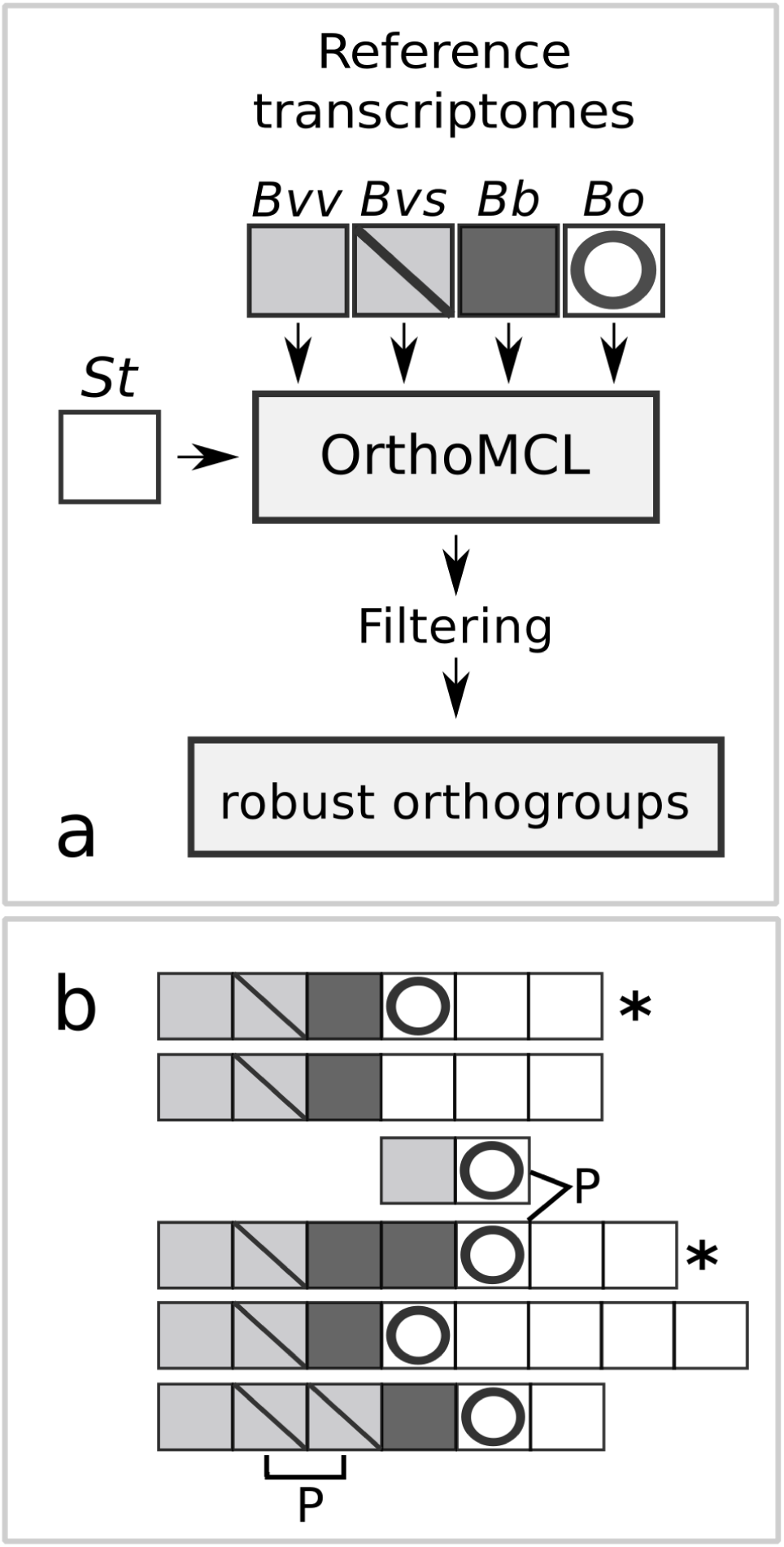
OrthoMCL clustering and filtering of raw groups. a. workflow. (taxon abbreviations as in Table 1, *St* = *Silurana tropicalis*) and b. examples of OrthoMCL clusters. Two clusters (indicated by an asterisk) pass the filtering criteria 1-3 (see text). Paralogue pairs (P) are linked by heavy lines.

### Variant detection

For the coalescence analyses, only the Illumina datasets were considered (corresponding to one diploid individual per taxon). For each taxon, reads were mapped to the reference transcriptome (for *B.b.* and B.v.v.: Trinity contigs only) with Bowtie2 v.2.2.4 (Langmead and Salzberg 2012, mode = sensitive). Variant detection was carried out with GATK (McKenna et al. 2010). Duplicates were removed, reads from isoforms were split as required (Split’N’Trim) and base qualities were recalibrated once to convergence. Raw variants were called with the HaplotpeCaller (genotyping mode = Discovery, stand_emit_conf = 10, stand_call_conf = 20) and filtered following recommendations for RNAseq data (DePristo et al. 2011; Van der Auwera et al. 2013).

Peptide alignments between taxon pairs within orthogroups were back-translated into the nucleotide sequences of the reference contigs with PAL2NAL (Suyama et al. 2006). For a given orthogroup and taxon, an alternate haplotype was constructed from the variant call format (VCF) file and added to the between-taxon alignments. These were parsed for variation at fourfold degenerate sites.

### Coalescent analyses

We used two complementary strategies to estimate models of divergence with and without gene flow analytically for each pairwise comparison of *B. v. variegata* against increasingly diverged taxa (*B. v. scabra, B. bombina, B. orientalis*). For an unrooted alignment of a pair of diploid samples, there are four types of sites: i) heterozygote in taxon A (k_A_), ii) heterozygote in taxon B (k_B_), iii) double heterozygote (2 distinct alleles only, k_AB_) and iv) alternate homozygotes (k_AABB_). First, we based inference on the average frequencies of these site types *E[k*_i_], essentially the folded, joint site frequency spectrum (SFS) for two species. Second, we used the joint distribution of the four site types in the short blocks of fixed length (150 consecutive fourfold degenerate sites) to fit models of divergence and gene flow. The configuration of mutations in each block can be summarized as a vector of site counts *<u>k</u> = {k*_A_, *k*_B_*,k*_AB_*,k*_AABB_*}*. Assuming blocks are unlinked, the overall model likelihood is given as the product of *p(<u>k</u>),* the probability of each observed blockwise configuration of mutations across blocks. We will refer to this as the blockwise SFS (bSFS) throughout.

These two approaches trade off power against bias. The SFS ignores all linkage information and summarizes the data into four site frequencies (Tab. 2). In principle, this allows us to co-estimate four parameters. However, the near absence of shared heterozygote sites, E[*k*_AB_], restricted our analyses to three-parameter models. This limitation is offset by more robust inference. In particular, point estimates based on the SFS are not affected by recombination or heterogeneity in mutation rate. In contrast, the analyses based on the bSFS assume a constant mutation rate across blocks and no recombination within them. Violations of these assumptions produce biased estimates. This is of particular concern for transcriptome data, given the added potential for recombination in introns. But by considering the joint distribution of linked mutations, the bSFS allows us to fit more complex models.

Lohse et *al.* (2015) have recently used a recursion for the generating function of genealogies (Lohse et al. 2011) to derive expressions for both *E[k*_i_] and *p(k)* under a model of isolation with unidirectional migration (IM). The model assumes an ancestral population of constant effective size N_anc_ that splits into two populations A and B at time *T* (measured in 2*N*_anc_ generations) followed by unidirectional migration at a constant rate of *M*=*4N*_anc_ *m* individuals per generation, where *m* is the proportion of immigrants per generation. We refer to the null model without migration as Div (Fig. 3)

**Figure 3.**
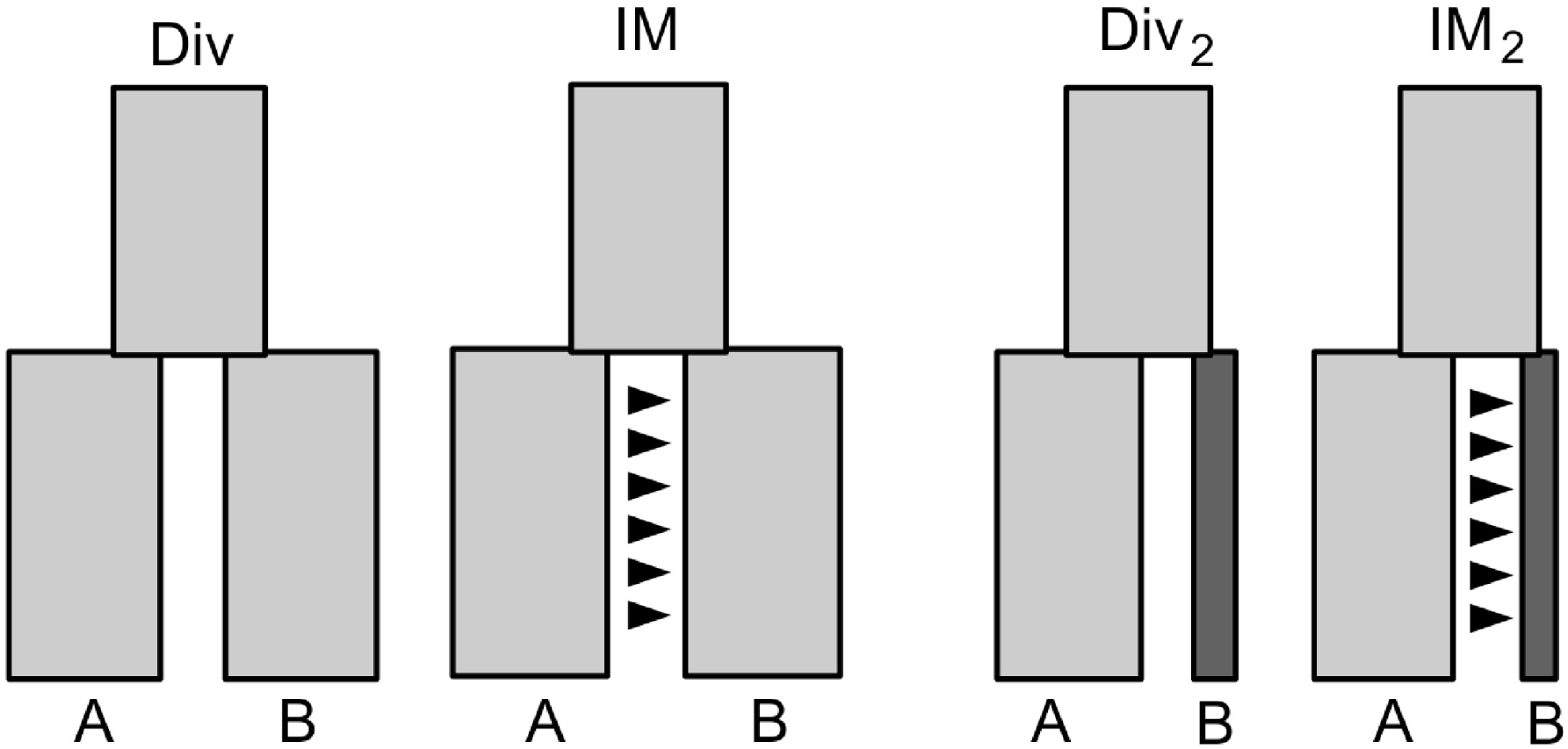
Sketches of the four coalescence models. The block widths indicate relative effective population sizes in the ancestral lineage and its two descendants A and B. Arrows mark the direction of gene flow.

Asymmetry in the frequency of unique heterozygous sites in each species (k_A_ and k_B_) may be due to differences in effective population size, asymmetry in post-divergence gene flow or both. We therefore also considered a model in which **N**_e_ in one of the descendant species was allowed to change instantaneously after divergence, while **N**_e_ of the other species remained constant (Fig. 3). We contrasted the relative support for this model without gene flow (Div_2_, **M** = 0) with that for its counterpart, IM, with gene flow but a single **N**_e_ for all three populations. These models with three parameters were fit using the SFS and the bSFS. Finally, we used the bSFS to fit models that allow for both post-divergence gene flow and asymmetries in **N**_e_ (**IM**_2_). In each case, we tested for gene-flow and asymmetries in **N**_e_ in either direction (i.e. *N*_A_ = a ***N***_anc_ or **N**_B_ = b *N*_anc_ and *M*_AB_ or *M*_BA_).

We assume that orthoblocks are randomly distributed and, given the large size of the *Bombina* genome (2600 cM, Morescalchi 1965), that the effects of physical linkage between them can be ignored (a rough calculation gives a maximum density of one orthoblock per 1.8 cM). Each orthoblock can therefore be treated as an independent replicate of the coalescent. To obtain a set of unlinked SNPs, we randomly sampled a single SNP per orthoblock and bootstrapped the analysis across 1000 such sub-sampled data sets. 95% confidence intervals of parameter estimates were estimated as 1.96 SD of parameter estimates across bootstrap replicates. The dataset comprised more than 1000 blocks with 150 fourfold degenerate sites and an average of 2.7 - 6.0 variable sites per block (depending on the comparison, Table 2). Maximum likelihood estimates were obtained using the *Nmaximise* function in *Mathematica* (for details see Lohse et al. 2015).

**Table 2.**
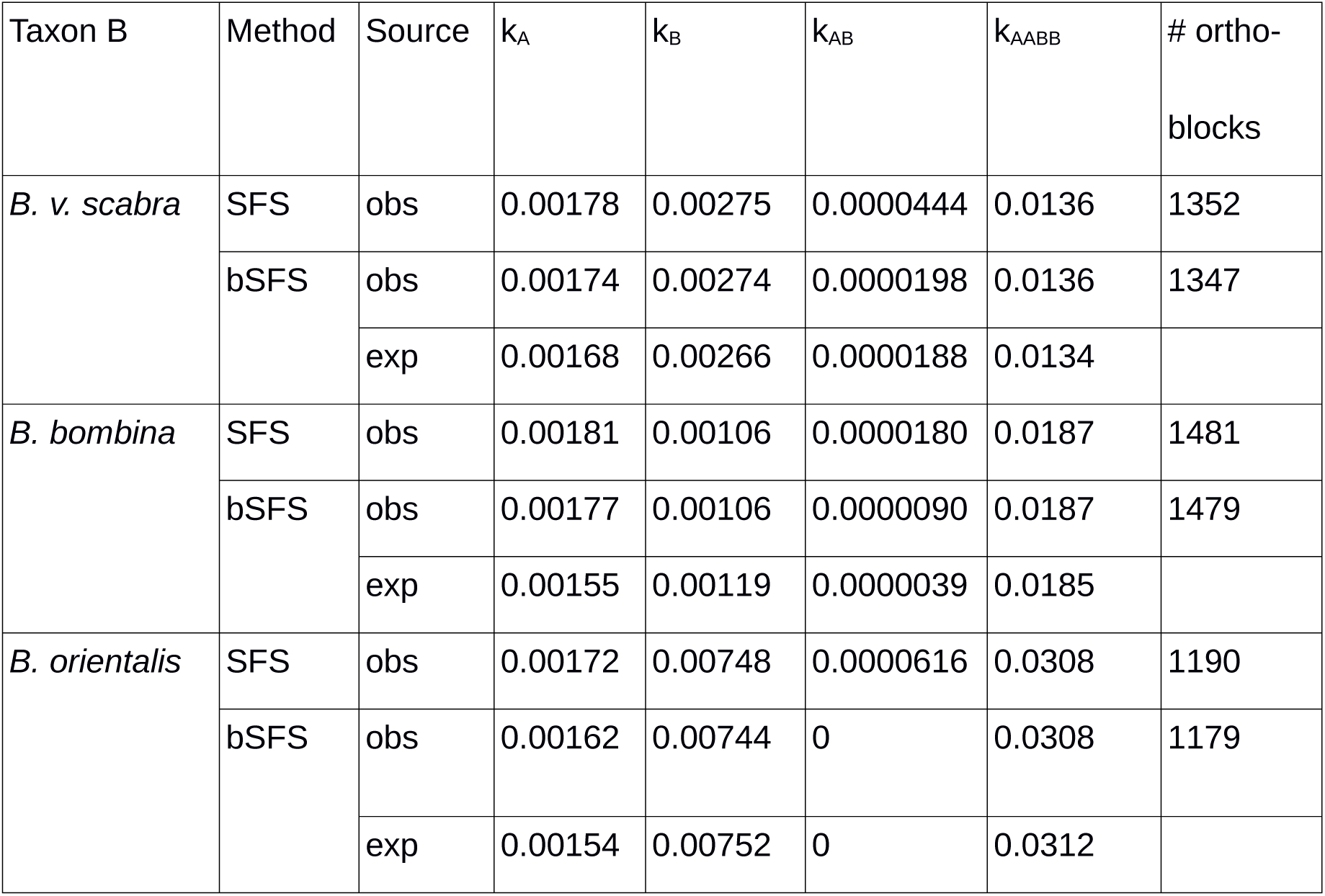
Average per site frequencies of the four mutation types for each pairwise comparison (A versus B). Taxon A is *B. v. variegata.* Observed data are listed separately for the site frequency spectrum (SFS) and the blockwise site frequency spectrum (bSFS). For the latter, the expected values from the best fitting model (Table 3, bold) are also given. The number of orthoblocks differs slightly between analyses, because all blocks that violated the four gamete test were excluded from the blockwise analysis.

| Taxon B | Method | Source | $k_A$ | $k_B$ | $k_{AB}$ | $k_{AABB}$ | # ortho-blocks |
| --- | --- | --- | --- | --- | --- | --- | --- |
| *B. v. scabra* | SFS | obs | 0.00178 | 0.00275 | 0.0000444 | 0.0136 | 1352 |
|  | bSFS | obs | 0.00174 | 0.00274 | 0.0000198 | 0.0136 | 1347 |
|  |  | exp | 0.00168 | 0.00266 | 0.0000188 | 0.0134 |  |
| *B. bombina* | SFS | obs | 0.00181 | 0.00106 | 0.0000180 | 0.0187 | 1481 |
|  | bSFS | obs | 0.00177 | 0.00106 | 0.0000090 | 0.0187 | 1479 |
|  |  | exp | 0.00155 | 0.00119 | 0.0000039 | 0.0185 |  |
| *B. orientalis* | SFS | obs | 0.00172 | 0.00748 | 0.0000616 | 0.0308 | 1190 |
|  | bSFS | obs | 0.00162 | 0.00744 | 0 | 0.0308 | 1179 |
|  |  | exp | 0.00154 | 0.00752 | 0 | 0.0312 |  |

**Table 3.** Support (ΔL) relative to the best model (*). Taxon comparisons are listed as in Table 2. SFS = site frequency spectrum, bSFS = blockwise analysis. Div = strict divergence; fixed N_e_, IM = divergence with migration, fixed N_e_; Div = strict divergence, asymmetric N_e_; IM_2_ = divergence with migration, asymmetric N_e_. Migration is unidirectional (A → B or B → A). See text for details. * The IM_2_ models collapsed to a strict divergence model (Div_2_)

| Taxon B | Method | Div | IM<sub>A→B</sub> | IM<sub>B→A</sub> | Div<sub>2</sub> | IM<sub>2,A→B</sub> | IM<sub>2,B→A</sub> |
| --- | --- | --- | --- | --- | --- | --- | --- |
| *B. v. scabra* | SFS | -30.2 | **0\*** | -13.2 | -22.4 |  |  |
|  | bSFS | -39.3 | -26.1 | -31.1 | -39.3 | -25.6 | **0\*** |
| *B. bombina* | SFS | -14.4 | -4.24 | **0\*** | -11.7 |  |  |
|  | bSFS | -58.0 | -26.3 | -0.3 | -43.6 | -3.0 | **0\*** |
| *B. orientalis* | SFS | -64.7 | **0\*** | -55.8 | -14.7 |  |  |
|  | bSFS | -223 | -223 | -223 | **0** | Div<sub>2</sub>* | Div<sub>2</sub>* |

### Estimation of synonymous substitution rates

In order to convert coalescence time estimates, *T*, into absolute times, *t*, we derived an estimate of the substitution rate per generation, μ, at fourfold degenerate sites from a recent anuran phylogeny based on nine nuclear genes (*rag1, rag2, bdnf, pomc, cxcr4, slc8a1, scl8a3, rho, h3a,* Irisarri et al. 2012b). That study demonstrated considerable rate variation between Neobatrachians and Archaeobatrachians (including *Bombina*). We therefore used data from two taxa in the latter group, *Alytes* and *Discoglossus* (MRCA 153 Mya, Roelants et al. 2011), which form a sister lineage to *Bombina.* From the concatenated nuclear dataset (Irisarri et al. 2012a), 896 fourfold degenerate sites were extracted and the substitution rate was estimated with PAML v. 4.7. (Yang 2007). Given the three year generation time of *Bombina,* we obtained μ = 9.15 × 10^−9^.

## Results

### Assembly overview

As is typical for Trinity assemblies, a large fraction (~ 60%) of relatively short contigs with low coverage had no similarity (BLASTX) with *S. tropicalis* and no ORF prediction. For example in the *B. v. variegata* assembly, contigs without ORF (n = 56,517) were shorter (median: 411 vs. 1214 *nt*) and had lower coverage (fpkm median: 1.14 vs. 2.19, mean: 3.76 vs. 27.3) than those with ORF prediction (n = 24,329). Similarly, in the Roche 454 assemblies, for *B. v. variegata,* contigs without ORF (n = 19,088) had a median length of 458 *nt* compared to 700 *nt* in contigs with ORF (n = 9739). The number of annotated Trinity components was close to the total gene count in other anurans (*S. tropicalis:* 26,800, *N. parkeri:* 23,400; Sun et al. 2015) and over 90% of 248 ultra-conserved core eukaryote genes (CEGs) were either partially or completely (> 70% of total protein length) assembled for each taxon.

### Paralogs

We focused our search for paralogs on those subsets of the assemblies in which ORFs had been identified, because all downstream analyses were based on coding sequence. In the Trinity assemblies, there were between 2791 (*Bvs*) and 3715 (*Bvv*) components per taxon with ORFs and more than one contig. Paralogs were identified in about 25% of these components in each taxon. In *B. v. variegata* for example, 10,753 contigs were considered in the paralog search and partitioned as follows: 3,715 contigs (with maximum ORF length) representing the components, 1,767 paralogs and a remainder of 5,271 contigs, that was removed from further analysis and presumably contained alleles and isoforms. For the other taxa, the paralog counts were 1,375 (*Bvs*), 1,737 (*Bb*) and 1,422 (*Bo*). A BLASTN search of Roche 454 contigs with ORFs against the Trinity database generated matches for 9,172 and 7,866 contigs in *B. v. variegata* and *B. bombina,* respectively. Of these Roche 454 contigs, 3,375 (Bvv) and 2,626 (Bb) were designated as paralogs. This relatively greater proportion reflects the overall lower identities of these BLAST-derived sequence pairs compared to those within Trinity components. Note that only the most similar Trinity/Roche 454 paralog pairs are expected to affect the definition of orthogroups (see the Supporting Information for additional information).

### Orthologs

Of the 15,775 OrthoMCL orthogroups with more than one *Bombina* taxon, 1,719 were excluded because they comprised at least one pair of paralogous contigs. There were 6,772 groups from which at least one of the four *Bombina* taxa was missing. An excess (> 3) of S. *tropicalis* contigs excluded 3,845 groups. *Bombina* paralogs were the sole reason for exclusion in 716 cases (Figure 4a). After these filtering steps, 4,978 orthogroups remained. Among these, 4,040 groups were robust to changes in the inflation value and used to generate pairwise alignments among taxa. The inclusion of presumptive paralogs in the clustering analysis not only led to the removal of groups with in-paralogs but also allowed for a sequence-based decision which transcript of a paralog pair to include in a given group. Figure 4b gives an example where paralog pairs were split into two orthogroups that most likely reflect alternate exon use in a gene with homology to acid phosphatase 1 in S. *tropicalis.* Note that in this case an alternative clustering based on just one contig per component resulted in an orthogroup that mixes isoforms and would lead to erroneous estimates of sequence divergence (see Supporting Information B for a comparison of both clustering approaches).

**Figure 4.**
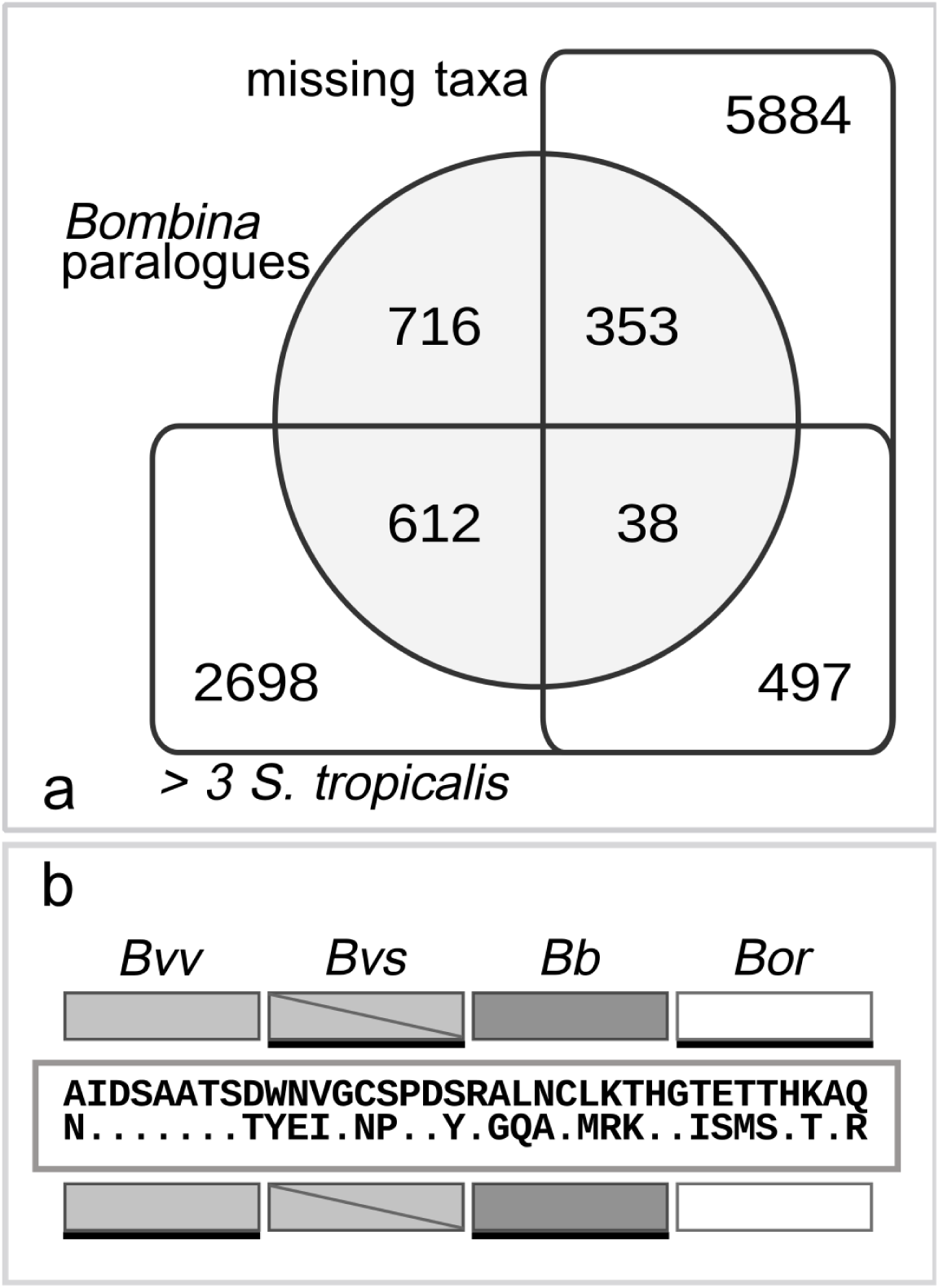
Filtering of OrthoMCL clusters. a. Counts of excluded clusters according to criteria 1 - 3 (see text for details). b. Examples of alternative exon use generating two orthogroups (rows of boxes), each with a member of a paralogue pair from each taxon. Shown are positions 41 - 76 (total: 158) of the aligned peptide sequence with differences between orthogroups (top *versus* bottom variant). A alternative set of reference transcriptomes comprising only the longest contig per component results in one cluster which mixes isoforms (heavy bars under the boxes).

### Variant detection

In each taxon, genotype qualities (a Phred-scaled measure of the confidence in the reported genotype) reached the maximum reported value (99) for more than 2/3 of SNPs and were 40 or over for over 95% of SNPs. We observed a small number of mostly low quality (< 20) genotype calls (0.5 - 3.4% per taxon) that did not include the reference allele (*i.e.* alternate homozygotes). With only one diploid individual per taxon, these cases point to assembly errors and were removed from further analysis.

Average heterozygosity (k_A_, k_B_) at fourfold degenerate sites ranged from 0.1 % in *B. bombina* to 0.7 % in *B. orientalis* (Table 2). Per site synonymous divergence (k_AABB_) was lowest between the subspecies *B. v. scabra* and *B. v. variegata,* intermediate between *B. variegata* and *B. bombina* and greatest between *B. bombina* and *B. orientalis* (Table 2). This is expected given the order of taxon divergence previously inferred from protein and mitochondrial data (Pabijan et al. 2013).

### Coalescence analysis

In both the *B. v. variegata/B. v. scabra* and the *B. variegata/B. bombina* comparison, an IM history fit the data significantly better than a history of isolation without gene flow. This was true regardless of whether we allowed for differences in **N**_e_ or not (Table 3) or whether the inferences was based on the SFS or the bSFS. Estimates of gene flow (*M*) were low (< 0.1) in both comparisons, but *M* was significantly greater between *B. v. variegata* and *B. v. scabra* than between *B. v. variegata* and *B. bombina,* both in the SFS and the bSFS (Table 4).

**Table 4.** Maximum likelihood estimates of parameters under the best supported model (Table 3). The divergence time (*T*) is given in units of 2 *N*_anc_ generations (95% CI in brackets). *a* is a scalar for the effective population size of one of the descendant populations (in parentheses, cf. Figure 3). n/a = not applicable.

| Taxon B | Method | $\theta$ | $M$ | $T$ | $a$ |
| --- | --- | --- | --- | --- | --- |
| *B. v. scabra* | SFS | 0.00179 | 0.078 (0.055, 0.099) | 12.5 (9.2, 15.7) | n/a |
|  | bSFS | 0.00129 | 0.032 | 12.0 | 2.07 (B) |
| *B. bombina* | SFS | 0.00107 | 0.027 (0.013, 0.041) | 22.4 (16.6, 28.1) | n/a |
|  | bSFS | 0.00121 | 0.020 | 16.8 | 0.99 (B) |
| *B. orientalis* | SFS | 0.00277 | 0.136 (0.120, 0.150) | ∞ | n/a |
|  | bSFS | 0.00752 | 0 | 3.75 | 0.2 (A) |

Our estimates of the divergence time also agreed well between SFS and bSFS analyses (Table 4). We converted estimates of *T* into absolute time using *t = T * 2N * g,* where is *g* is generation time, and *N = θ/(4μ),* using an estimate of *μ* from a related pair of Archaeobatrachians (*cf.* Methods). Based on the bSFS analysis, this resulted in ancestral population sizes of 0.33 - 2 × 10^5^. The divergence times for *B. v. variegata* and *B. v. scabra* were 3.6 and 2.5 mya for the SFS and the bSFS analysis, respectively. The analogous figures for the other comparisons were 3.9 and 3.3 mya (*Bvv/Bb*) and 4.6 mya (*Bvv/Bo*).

The SFS and bSFS analyses differed in several respects. First, the blockwise analyses showed no support for gene flow in the allopatric comparison of *B. variegata* and *B. orientalis.* In other words, allowing for gene flow as well as differing population sizes (IM_2_) did not improve the fit compared to a scenario of strict divergence with varying N_e_ (Div_2_, Table 3). In contrast, the SFS approach favored a model of strong gene flow (M = 0.136) from *B. v. variegata* to *B. orientalis* (IM) over Div_2_. Second, using the SFS the best fitting model involved gene flow and always into the gene pool with relatively greater heterozygosity (*e.g.* from *B. bombina* to *B. v. variegata*). In contrast, the bSFS analyses could separately account for varying N_e_ and, in the case of the two *B. variegata* subspecies, gave significantly better support for gene flow in the opposite direction (*i.e.* from *B. v. scabra* to *B. v. variegata).* Finally, estimates of *M* were lower in the bSFS than the SFS analyses (Table 4).

To assess the relative fit of the best supported models, we computed the expected SFS under the models inferred from the bSFS (Table 2). We also tested how much of the gene flow signal is due to the presence of shared heterozygous sites. For *B. variegata* and *B. bombina,* only one block contained shared heterozygous sites and removing this block had no effect on the results. Instead, most of the signal for gene flow came from an excess of blocks with no fixed differences (k_AABB_= 0, Fig. 5). For *B. variegata* and *B. bombina* there are 134 such blocks which is highly unlikely under the best fitting divergence scenario (p= 0.002 given a total of 1,479 blocks and assuming binomial sampling), but not under the IM_2_ model (p = 0.89). We can use analogous coalescent calculations to ask whether there is an excess of invariant blocks (<u>k</u> = {0,0,0,0}), as would perhaps be expected if introgression between *B. variegata* and *B. bombina* occurred in a recent burst following the formation of the hybrid zone, or was partially adaptive. There were 95 invariant blocks, which is not unexpected under the IM_2_ model (p = 0.235).

**Figure 5.**
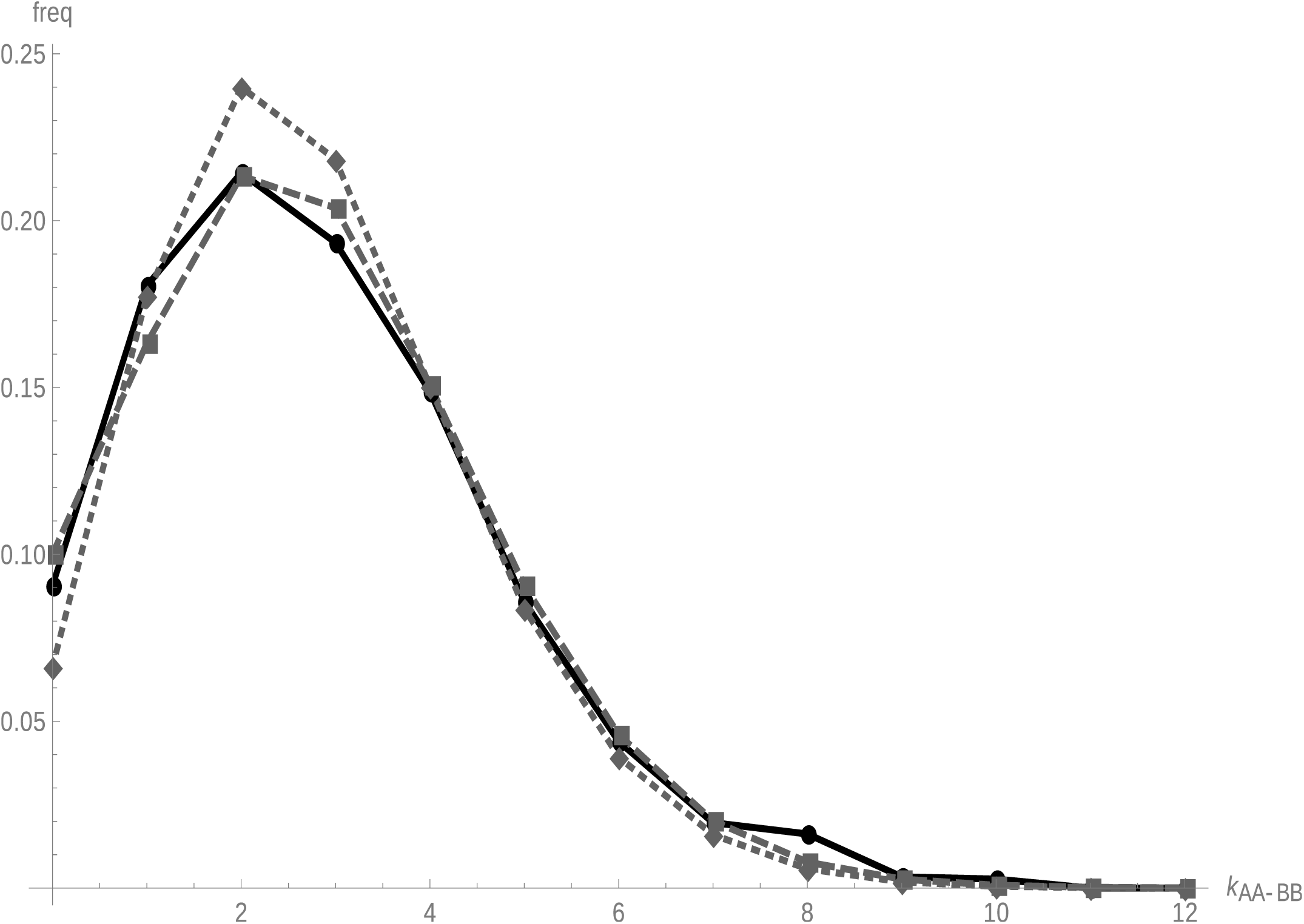
The distribution of fixed differences between *B. v. variegata* and *B. bombina* (k_AABB_) in blocks of 150 fourfold degenerate sites. The divergence model predicts a narrower distribution (gray dotted) than observed (black), *i.e.* there is an excess both of blocks with no fixed differences and of blocks with a large number of fixed differences (k_AABB_ > 6). An IM_2_ model with migration and asymmetry in N_e_ (gray, dashed) gives a good fit to the data.

## Discussion

The coalescent analyses provide evidence that hybridization between *B. bombina* and *B. v. variegata* results in continued gene exchange between these anciently diverged taxa, but the estimated long term average of gene flow was very low (M = 0.020 or 0.027, depending on the analysis). Thus despite the abundance of fertile hybrids in contact zones, the effective rate of gene exchange between the two pure populations was no greater than one immigrant per 100 - 150 years. These values of M are at least 40 times smaller than an estimate by the same method for two sympatric *Heliconius* species that hybridize infrequently (Lohse et al. 2015). However when integrated over time, the *Bombina* figures imply that a substantial portion of genes has been affected by past gene flow: based on the results from the bSFS analysis, this proportion for *B. v. variegata* is 1 – e^−0 020^ * ^168^ = 0.285 (Table 4). The analogous figure for *B. bombina* is 0.24.

Before we explore our results in more detail, we comment on aspects of our methodology. We devised an analysis pipeline that identified putative paralogs based on nucleotide sequence identity and excluded all cross-taxon gene clusters with paralog pairs from further analysis. This filtering step will also have eliminated the more serious cases of incorrectly phased variants (Haas et al. 2013). By using the same tissue (liver) for all samples, we avoided problems due to tissue-specific expression. However, some errors will remain. For any recent paralog pairs above our identity threshold (0.98), only one contig was used in the clustering step and so orthology may have been mis-assigned. Finally, differential gene loss (Rasmussen and Kellis 2012) may have resulted in some paralogous clusters.

Interpretation of the coalescent results requires weighing the strengths and weaknesses of both inference schemes. Recombination within blocks reduces the variance in coalescence times in the bSFS scheme and so leads to underestimates of *M* and *N*_e_ (Wall 2003). Our blocks of 150 fourfold degenerate sites were on average 1500 nt long, which is longer than the average CDS length in *S. tropicalis* (1323 nt) and *N. parkeri* (1382 nt, Sun et al. 2015). In these fully sequenced genomes, the average gene length is 18.0 kb and 24.4 kb, respectively (Sun et al. 2015). This suggests an average span of 20 kb or more for the genomic complements of our blocks and a likely influence of recombination on the parameter estimates (recall the relatively lower values for *M*, Table 4). Overall, though, the biases appear small, because the estimates for a given model agreed well between bSFS and SFS analyses. Selective sweeps at linked sites in the ancestral population may have the opposite effect of inducing additional variation in coalescence times (Coop and Ralph 2012). The excess of blocks with no or few divergent sites in the European comparisons may in principle be explained in this way. But no such excess is seen in the allopatric comparison (*Bvv/Bo*) which suggests that it is instead a genuine signal of past migration.

While the SFS analysis makes fewer assumptions, it confounds gene flow with differences in effective population size since the lineage split. Thus, when assuming a single *N*_e_, we always inferred gene flow into the taxon with higher heterozygosity and obtained unrealistic estimates for the allopatric pair (Table 4). In the bSFS analyses with two *N*_e_ parameters, *M* estimates were reduced to zero as expected (*Bvv/Bo*) or the direction of gene flow was reversed (*Bvv/Bvs*). The bSFS contains additional information to distinguish gene flow from asymmetries in *N*_e_: a larger *N*_e_ in one descendant population increases heterozygosity uniformly whereas gene flow generates a small number of blocks with large numbers of heterozygous sites in the population receiving migrants.

The observed heterozygosities per taxon (Table 2) fully match our expectations from previous studies. *B. bombina,* with the lowest estimate (k_B_ = 0.00106), has recently expanded its range from glacial refugia near the Black Sea and has displayed minimal genetic diversity in all previous *Bombina* comparisons (Szymura 1988, 1993; Fijarczyk et al. 2011). The difference in heterozygosity between the *B. variegata* subspecies agrees with previously reported differences in *N*_e_ (Fijarczyk et al. 2011). Finally, the *B. orientalis* specimen (max. estimate in this study, k_B_ = 0.00748) comes from Korea, an important Pleistocene refugium in Asia (Zheng et al. 2009).

While the blockwise analyses allow a more detailed reconstruction of the divergence history, they still fit a very simplistic model. In particular, the assumption of unidirectional gene flow is unlikely to apply to *Bombina.* In fact for the hybridizing taxa, both the support for the two versions of the IM_2_ model (A→B vs. B→A, Table 3) and the M estimates under these models are very similar (M in the less favored direction (*Bvv→Bb*) is 0.0127). Note also that the support for asymmetric N_e_ is very weak and M for the IM model with a single N_e_ (*Bb →Bvv*) is also 0.02. For the two *B. variegata* subspecies, the best supported model (IM_2,b→a_) matches previous findings of relatively more gene flow from the Balkans into the Carpathians (Fijarczyk et al. 2011). Given these limitations and considerations of bias above, *M* estimates based on the bSFS should be interpreted as a minimum, long-term average of gene flow following the lineage split.

The species divergence times we have inferred here are by necessity more recent than the divergence at any particular gene (which includes an additional coalescence time in the common ancestral population). However, the difference between our estimates and previous calibrations for mitochondrial genes is surprising. For example for the split between *B. v. variegata* and *B. bombina,* we obtain 3.3 Mya compared to the 6.4 or 8.9 Mya divergence at mtDNA genes (two different fossil calibrations, Pabijan et al. 2013; see also Heled and Drummond 2015). While some of this difference may reflect coalescent stochasticity (*i.e.* by chance, the coalescence of mitochondrial genomes may considerably predate the taxon split), it likely reflects differences in calibration.

The new divergence time estimate is consistent with the appearance of the first *Bombina bombina* fossils in the late Pliocene (Rage and Rocĕk 2003) and with analyses of allozymes (Nei’s D, Szymura 1993). But it too is clearly beset by uncertainties. We therefore conclude cautiously that *B. bombina* and *B. v. variegata* diverged in the mid-Pliocene, at a time when much of their present-day distribution range in Central Europe was still covered by remnants of the Paratethys Sea. Subsequent climatic oscillations during the Quarternary will have resulted in repeated range shifts (*cf.* Futuyma 1987), presumably with recurring opportunities for gene exchange that have left a clear signal in present-day transcriptomes.

However, our dataset contains no signal of increased introgression due to ongoing hybridization. While the proportion of invariant blocks is significantly larger than expected under strict divergence, it is consistent with a constant rate of gene flow. This observation agrees with the prediction from cline theory (cf. Introduction): considering the barrier effect (51 km) and the distance of the *B. v. variegata* sample from the hybrid zone (61 km), it should take roughly 18,500 years for genetic variants to cross the Kraków transect and arrive at Biała Woda.

Assuming similar barrier strengths, the *B. bombina* sample (71 km) should be insulated even longer from introgression *via* that route. Selectively favored alleles, however, will introgress much more quickly (Barton 1979). For example, an allele with a 1% selective advantage is expected to traverse the Kraków transect in a mere 245 years (*via* expression in Barton and Gale, 1993, p. 30 top). Our data set may well contain a few blocks of this kind, but larger samples of individuals and long range linkage information would be needed to detect them.

The strength of the genetic barrier to gene flow varies among different *Bombina* contact zones. Near Apahida, Romania, the two taxa meet in an extended habitat mosaic at a distance of 20 km from the nearest pure species range (Vines et al. 2003). Plausible total selection strengths can be found that could balance the high estimated gene flow rates and maintain adaptive taxon differences indefinitely. But with, say, 20 selected loci, neutral marker clines would dissipate within ~ 160 years. A very dense spacing of the total selection pressure would be required (~ one selected locus every 4 cM, *i.e.* about 650 loci) to maintain such clines over 10,000 years. In fact, there are reasons to believe that this particular contact zone is much more recent (Vines et al. 2003). Spatial and temporal heterogeneity in rates of gene exchange is a common theme in the hybrid zone literature (Harrison and Larson 2014) and often linked to environmental conditions, as in this example. Episodic spates of gene flow appear to be just as much a product of environmental heterogeneity as any initial adaptive divergence. But given the slow rate of diffusion, neutral introgression should attenuate quickly with increasing distance from the hybrid zone, even in Apahida.

A post-zygotic barrier to neutral gene flow depends on physical linkage between neutral and selected loci and thus becomes stronger as a given total selection stength is more finely distributed across the genome (Barton and Bengtsson 1986). In *Bombina,* the number of loci under selection has been estimated to be at least 55 and is most likely considerably larger (Szymura and Barton 1991). Abundant trait differences also suggests that multifarious selection may promote reproductive isolation, not just because many loci are involved but also because of the pleiotropic interactions they may have (Nosil and Harmon 2009). Under Fisher’s (1930) geometric model, such interactions lengthen the adaptive walk towards a new phenotypic optimum and, as a consequence, reduce the mean fitness of hybrids upon secondary contact (Chevin et al. 2014). The largest drop in mean hybrid fitness is expected after the first few mutations have fixed, but the total fitness reduction at the end of an adaptive walk is only moderate (< 50%, Barton 2001). This polygenic model of adaptive divergence thus predicts that hybrids would continue to be conduits of gene flow unless adaptive substitutions also trigger strongly deleterious epistatic interactions (i.e. Dobzhansky-Muller incompatibilities, Chevin et al. 2014). Such incompatibilities exist in *Bombina (cf.* Introduction). They may or may not result from ecological divergence, but their joint effect is clearly insufficient to cause reproductive isolation. We do not know at present why this is so but note that one important driver of speciation is missing: these toads do not have heteromorphic sex chromosomes and so Haldane’s Rule (Orr 1997) does not apply.

As it stands, *B. bombina* and *B. variegata* play important roles in their respective ecological communities, just like ‘good’ biological species, despite their gene pools fraying at the edges. With an estimate of *m* on the order of 10^−6^ only a few dozen dispersal ranges away from the hybrid zone, adaptive divergence can proceed uninhibited by hybridization. Similarly low gene flow rates were obtained for hybridizing sunflower species (Sambatti et al. 2012). Under these para-allopatric conditions (Coyne and Orr 2004), there is no antagonism between selection and gene flow. Consequently, any speciation mechanism may increase reproductive isolation between well separated, pure population of either taxon without constraints on the underlying genetic architecture. In contrast, local adaptation under selection-gene flow balance between two panmictic populations favors a clustered genomic distribution of selected loci (Felsenstein 1981; Ortíz-Barrientos et al. 2002) which can gradually evolve in redundant genetic systems (Yeaman and Whitlock 2011). Especially when such traits under divergent natural selection also cause assortative mating (magic traits, Gavrilets 2004; Servedio et al. 2011), novel ecotypes may rapidly form. But as long as prezygotic barriers are incomplete or labile under environmental perturbations, hybrids will be generated and post-zygotic barriers to gene flow are likely to be inefficient.

The conundrum of incomplete speciation in *Bombina* illustrates that speciation tends to defy rules (Coyne and Orr 1989) and argues for an open-minded exploration of possible paths towards reproductive isolation. Rapid evolution of reproductive isolation between recent ecotypes is feasible whenever it can overcome the challenges of a limited gene pool from which to draw mutations that drastically reduce gene flow, the trend towards a clustered genomic distribution of selected traits and the need to build up irreversible barriers to gene flow before further environmental change threatens to reverse diversification. This suggests that many young ecotypes will be ephemeral, because they do not persist long enough to reach complete isolation (Futuyma 1987). Those ecotypes that adapt to abundant and widely distributed resource should stand a relatively much better chance to become reproductively isolated in their own time. This does not diminish the role of ecology in generating biological diversity. To the contrary, it underscores that relatively young ecotypes can play important roles in their respective communities and even prompt diversification in other taxa (Abrahamson and Blair 2007), irrespective of whether they are reproductively isolated. We hasten to add that this view has a very long history indeed (Darwin 1859; see Mallet 2008).

## Data archiving

Raw Illumina reads were submitted to SRA with accession numbers PRJEB7348-PRJEB7351. *De novo* assemblies of Roche 454 and Illumina data including orthology information are available on the AfterParty platform (link). The source code for the paralog filter written in Perl can be found at github/…. (see note in Supporting Information A).

## Acknowledgments

We thank N. Wrobel (Edinburgh Genomics) and M. Kuenzli (Functional Genomics Center, Zürich) for help with RNA extraction and Roche 454 sequencing, respectively. W. Babik, B. Elsworth, M. Jones, G. Koutsovoulos, S. Kumar and D. Laetsch generously shared their bioinformatics and computing expertise. We thank N. Barton for comments on a previous version of this manuscript. BN was supported by a Daphne Jackson Fellowship funded by the Natural Environment Research Council. AF received funds from the European Science Foundation (FroSpects) and Society-Environment-Technology (www.set.uj.edu.pl) for an exchange visit in Edinburgh. Collecting permits were issued by the Minister of the Environment, Republic of Poland, DLKOPiK-op/ogiz-4200/II-6/1441/07/msz.s. The study was approved by the First Local Ethical Committee on Animal Testing, Jagiellonian University, Kraków (10/OP/2006). We acknowledge financial support from the Polish Ministry of Science and Education (N303 11832 / 2735) and the National Science Centre (2013/09/B/NZ8/03349) to JMS.

